# LAT condensation gates PLCγ1 activation via bimodal LAT phosphorylation

**DOI:** 10.64898/2026.04.29.721478

**Authors:** He Ren, Hyun-Ro Lee, Yannick A.D. Omar, Joseph B. DeGrandchamp, Chien-Lun Hung, Timothy J. Eisen, Dimitrios Stamou, John Kuriyan, Arup K. Chakraborty, Jay T. Groves

## Abstract

T cells can respond to even a single molecular binding event of antigen to a TCR. A key step in the TCR signaling pathway that definitively exhibits this single molecule response is the initiation of calcium influx by activation of PLCγ1 in the LAT protein condensate. Here, we describe detailed kinetic measurements examining how protein condensation of LAT regulates activation of PLCγ1 using a reconstituted membrane system. The results reveal that membrane recruitment of PLCγ1 is tightly controlled by the LAT phosphorylation state, with no measurable independent recruitment to PIP_2_ or PIP_3_ lipids via the PLCγ1 PH domains. We further observe PLCγ1 is rapidly activated by membrane-associated kinase upon recruitment, irrespective of the LAT condensation state. These studies also revealed a crosstalk mechanism in which the TEC family kinases responsible for PLCγ1 activation also phosphorylate LAT. This interaction establishes a positive feedback loop in LAT phosphorylation, mediated through LAT condensation, which drives a bimodal LAT phosphorylation response to TCR activation. Kinetic modeling reveals how this LAT phosphorylation response can cooperatively gate PLCγ1 activation from a single TCR. These results suggest the LAT condensate facilitates both signal amplification and noise suppression in PLCγ1 activation through a bimodal switch affecting LAT phosphorylation.

**Significance Statement:** T cells are sensitive sensors capable of detecting and responding to trace amounts of foreign antigen. Understanding how they achieve such sensitivity while maintaining accurate antigen discrimination remains a key challenge. Here, through detailed kinetic measurements of PLCγ1 activation, we identify a cross reactivity in which kinases responsible for PLCγ1 phosphorylation also phosphorylate LAT. This creates a bimodal switch controlling LAT phosphorylation levels, which gates PLCγ1 activation from single TCR signals. We suggest this mechanism plays a key role in the signal amplification and noise suppression required for T cells to detect single antigen molecules.

## Introduction

The T cell receptor (TCR) signaling pathway is acutely sensitive to antigenic stimulation and can respond even to a single molecular TCR activation event (1-3). Although the basic reaction sequence in TCR signaling is well mapped (4), a mechanistic understanding of its single molecule sensitivity is still limited. Peptide-major histocompatibility complex (pMHC) molecules on antigen-presenting cell (APC) surfaces that bind TCR for at least a few seconds lead to ITAM phosphorylation on the TCR followed by recruitment and activation of the Syk family kinase, ZAP70. ZAP70 has distinctive substrate specificity and phosphorylates the membrane-linked scaffold protein linker for activation of T cells (LAT) (5, 6). Activation of ZAP70 on a TCR does not guarantee downstream signal propagation (7, 8), indicating the presence of an as yet poorly resolved signal discretization step that gates productive signaling to LAT (9, 10). Once phosphorylated, pLAT then recruits a variety of downstream signaling proteins, including phospholipase C gamma 1 (PLCγ1), Grb2, the Ras activator Son of Sevenless (SOS), Gads, and SLP76 (11-14). These molecules become crosslinked in a multivalent bond percolation network through a cooperative process that has been called a form of phase transition (15-18) (Figure 1A).

**Figure 1.**
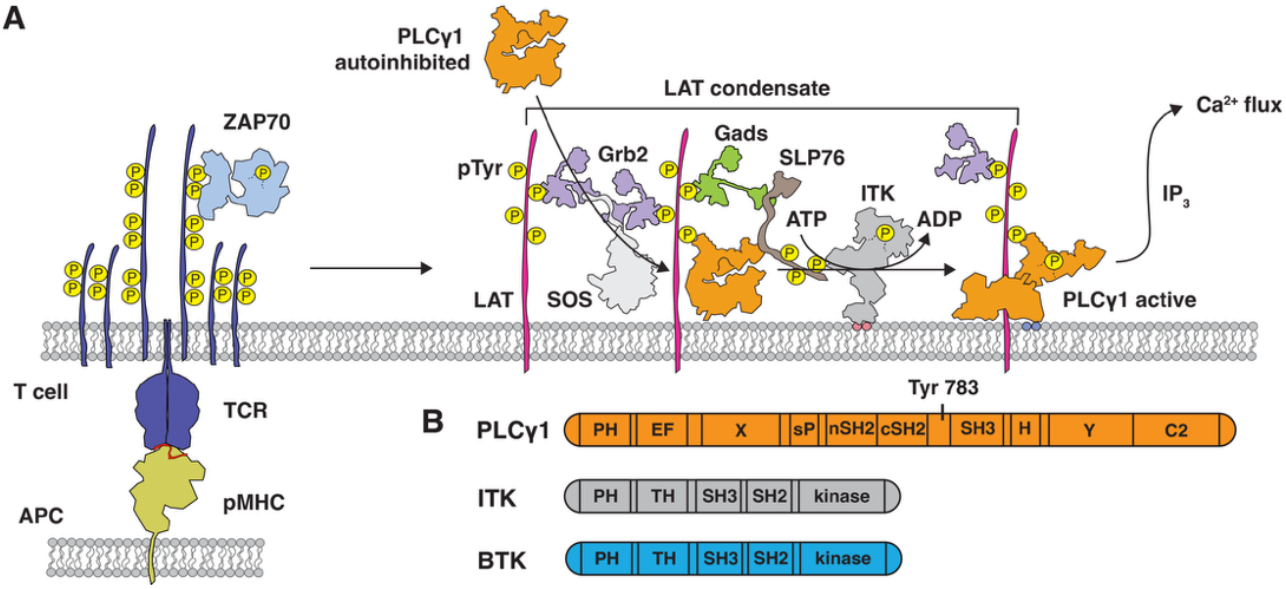
Schematic of the PLCγ1 activation assay. (A) T cell receptor engagement by antigen peptide-MHC initiates a kinase cascade that activates ZAP70 and phosphorylates LAT. Phosphorylated LAT (pLAT) recruits and gets crosslinked by multiple downstream signaling proteins through multivalent interactions. Upon formation of a mature LAT condensate, PLCγ1 becomes activated and triggers a cell-wide calcium spike. (B) Domain organization of PLCγ1, ITK, and the ITK surrogate BTK.

In living primary T cells, discrete LAT condensates have been observed to form in response to individual TCR activation events (9, 19). The condensation process is abrupt and occurs after a distinct delay time (averaging tens of seconds). Local pLAT levels near an activated TCR exhibit a correspondingly bimodal response, with stable pLAT buildup only observed in the condensate. Within seconds of formation, each LAT condensate contributes a single, cell-wide calcium spike (20), and this calcium influx marks one of the earliest indicators of T cell activation (21). These recent observations identify the LAT condensation process itself as a key player in signal gating and amplification to achieve single molecule sensitivity in TCR signaling.

While there is abundant speculation that condensates of signaling molecules are positive promoters of signaling (22-28), quantitative measurements or mechanistic understanding of this are limited. Activation of Ras by the Ras guanine nucleotide exchange factor, SOS, both in LAT condensates as well as in similar EGFR condensates, is an exception: SOS autoinhibition release and subsequent Ras activation is strongly promoted by the condensed state through a type of kinetic proofreading mechanism (29-32). PLCγ1 is also evidently activated rapidly in LAT condensates since it is the primary driver of calcium flux in T cells (33), and this calcium response is observed to occur nearly synchronously with LAT condensation (20). However, recent membrane reconstitution studies of PLCγ1 activity reveal PLCγ1 is highly active when recruited to dispersed LAT and does not depend on the LAT condensation state (34). The mechanism by which LAT condensates gate PLCγ1 activity is clearly distinct from SOS, even though they are both signaling from the same condensate. Here we describe a detailed kinetic examination of PLCγ1 activation in LAT condensates. The results lead to a different mechanism of condensate-mediated regulation rooted in positive feedback and bimodality in the kinase-phosphatase competitive reactions governing LAT phosphorylation itself.

PLCγ1 is a ubiquitously expressed phospholipase that hydrolyzes phosphatidylinositol 4,5-bisphosphate (PIP_2_) into inositol trisphosphate (IP_3_) and diacylglycerol (DAG). IP_3_ binds to IP_3_ receptors on the endoplasmic reticulum (ER) membrane, triggering the release of calcium ions into the cytoplasm. DAG recruits RasGRP and Protein Kinase C (PKC) to the membrane, contributing to Ras activation and other signaling processes

(35). PLCγ1 consists of a catalytic core along with regulatory domains that maintain it in an autoinhibited state prior to activation at the membrane. The regulatory domains—including a split PH domain, tandem SH2 domains (nSH2 and cSH2), and an SH3 domain—insert between the X and Y boxes of the catalytic core (Figure 1B). Recent crystal and cryo-EM structures have shown that these regulatory domains sit atop the catalytic core, blocking substrate access to the catalytic site (36-39). Phosphorylation of PLCγ1 at tyrosine 783 induces an intramolecular interaction with the cSH2 domain, resulting in a global conformational change that relieves autoinhibition (40, 41).

In T cells, PLCγ1 is primarily recruited to the membrane via its nSH2 domain binding to phosphorylated tyrosine 132 (pY132) on human LAT (pY136 in mouse) (42). In addition to LAT-dependent recruitment, PLCγ1 has also been reported to associate with membranes through direct interactions with phosphoinositide lipids, especially phosphatidylinositol 3,4,5-trisphosphate (PIP_3_) (38, 43, 44), but the contribution of these lipids to the PLCγ1 membrane recruitment remains unclear. Phosphorylated LAT also recruits Gads and SLP76, which in turn bring in the TEC-family kinase ITK, which phosphorylates tyrosine 783 on PLCγ1 to activate it (45-48). The SH2 and SH3 domain array of PLCγ1 further allows it to crosslink with multiple LAT molecules and facilitate the formation of protein condensates (49, 50). PLCγ1 activation is subject to tight regulation in T cells and its misregulation can lead to disease. In adult T cell lymphoma (ATL), for example, PLCγ1 is the most frequently mutated gene (51). Many of these mutations disrupt the regulation of PLCγ1 membrane recruitment and activation to contribute to disease progression (36, 38, 52).

In this study, we performed detailed kinetic analyses to elucidate the activation mechanism of PLCγ1 using a supported membrane reconstitution platform. This assay mimics the native signaling environment by integrating core regulatory events including receptor-mediated recruitment of autoinhibited PLCγ1 to the membrane, TEC family kinase-mediated phosphorylation of PLCγ1 after membrane binding (48), and PLCγ1-catalysis of PIP_2_ to generate DAG and IP_3_. Another advantage of the supported membrane configuration, in contrast to liposome- or solution-based assays (38, 53-55), is the ability to control the condensation state of LAT on the membrane (16, 29). Using total internal reflection fluorescence (TIRF) microscopy, we quantitatively measured the kinetics of both membrane recruitment and catalytic activity of PLCγ1, independently. The results indicate that PLCγ1 membrane recruitment is primarily governed by LAT phosphorylation, while its PH domain only engages with PIP_2_ and PIP_3_ lipids after pLAT-mediated recruitment. The mean binding dwell time of the pLAT-PLCγ1 interaction is two orders of magnitude longer than that of the pLAT-Grb2, identifying PLCγ1 as a major stabilizing element in LAT condensates. Following recruitment, autoinhibited PLCγ1 is rapidly activated by kinase on the membrane irrespective of LAT condensation state. These experiments also revealed an unexpected crosstalk potential between the TEC family kinase ITK and LAT.

Although ITK is primarily considered to be a PLCγ1 activator, we observe evidence that its kinase domain also phosphorylates LAT at rates that are significant in the condensate. This introduces a strong positive feedback loop that facilitates LAT condensate growth. Moreover, any ITK-driven LAT phosphorylation is independent of ZAP70 once the condensate is nucleated. Kinetic modeling confirms that this feedback can establish bimodality in the kinase-phosphatase kinetic competition governing LAT phosphorylation levels in T cell. Taken together, these findings suggest that the condensation phase transition of LAT establishes a bimodal switch governing LAT phosphorylation, whereby LAT remains phosphorylated within condensates but is rapidly dephosphorylated outside. In this way, LAT condensation regulates PLCγ1 activity not through structural features of the condensed phase, but by imposing tight spatiotemporal control over PLCγ1 recruitment and activation via bimodal LAT phosphorylation.

## Results

### PLCγ1 membrane recruitment requires pLAT

We first developed an in vitro reconstitution system to quantify the recruitment kinetics of PLCγ1 to the membrane, which is the initial step for PLCγ1 activation. Controlling the LAT density, LAT phosphorylation level, and PIP lipid composition in the membrane, we measured PLCγ1 recruited to the membrane over time (Figure 2A). Full-length PLCγ1 with an C-terminal mNeonGreen tag was transiently expressed in HEK293T cells and harvested as whole-cell lysate. The solution concentration of PLCγ1 in lysate was calibrated against purified mNeonGreen-tagged protein by epifluorescence microscope (Figure S1A). Supported lipid bilayers were prepared with DOPC, Ni-NTA-DOGS, with and without phosphoinositide lipids. The cytosolic domain of LAT, with an N-terminal His10 tag, was tethered to the bilayer via nickel-histidine interactions to achieve a surface density of approximately 500 molecules per μm^2^. LAT was pre-phosphorylated using the kinase domain of AncSZ, an ancestrally reconstructed Syk-family kinase (56), and separated by chromatography from the kinase prior to membrane incubation. The membrane density of PLCγ1 was calibrated in Figure S1B.

**Figure 2.**
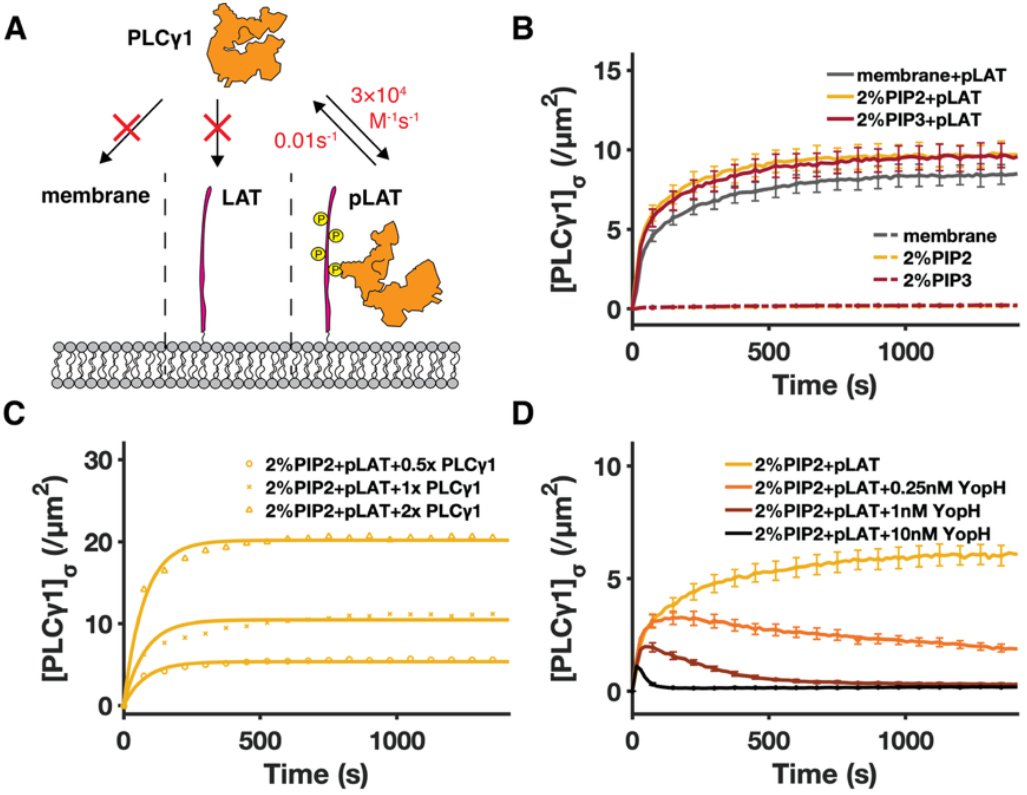
PLCγ1 membrane recruitment requires pLAT. (A) Schematic of the PLCγ1 membrane-recruitment assay. (B) PLCγ1 membrane recruitment requires pLAT, and direct PLCγ1-lipid interactions contribute minimally to membrane recruitment. Effects of phosphoinositide lipids emerge only after initial pLAT binding. pLAT was incubated at approximately 500 molecules/μm^2^, and PLCγ1 (1x) was added at 15 nM. […]_σ_ denotes surface density. Mean ± SD of 25 membrane patches. (C) PLCγ1 membrane-binding kinetics extracted from a PLCγ1 concentration titration. Dotted lines represent experimental recruitment traces, while solid lines represent simulations of a reversible pLAT-PLCγ1 binding model using globally fitted *k*_*on*_ = 3 × 10^4^ M^−1^s^−1^ and *k*_*off*_ = 1 × 10^−2^ s^−1^. (D) The pLAT-PLCγ1 interaction is reversible upon addition of the tyrosine phosphatase YopH. (membrane: 96:4 DOPC:Ni-NTA; 2% PIP_2_: 94:4:2 DOPC:Ni-NTA:PIP_2_; 2% PIP_3_: 94:4:2 DOPC:Ni-NTA:PIP_3_)

The results showed that PLCγ1 is specifically recruited to pLAT in membranes, with minimal direct recruitment to membranes via PIP_2_ and PIP_3_ (Figure 2B). Although both PIP_2_ and PIP_3_ moderately enhanced PLCγ1 recruitment on the pLAT-containing membrane, likely through nonspecific electrostatic interactions (38), their contribution to the PLCγ1 membrane recruitment was insignificant.

We examined the kinetics of pLAT-mediated recruitment of PLCγ1 by titrating the enzyme concentration in the recruitment assay (Figure 2C) and analyzing the data using a simple model of binding kinetics (See SI for details). These results indicate that PLCγ1 exhibits a relatively long membrane dwell time bound to pLAT (∼100 s). We further examined PLCγ1 recruitment to pLAT in the presence of varying amounts of YopH, a tyrosine phosphatase that dephosphorylates LAT (16). Upon simultaneous addition of YopH and PLCγ1, PLCγ1 initially localized to the membrane but gradually dissociated over time, with higher phosphatase concentrations accelerating the loss of membrane association (Figure 2D). The timescale of phosphatase activity is consistent with PLCγ1 unbinding kinetics inferred from the PLCγ1 concentration titration. The ∼100 s mean binding dwell time of PLCγ1 to pLAT contrasts that of Grb2, which is orders of magnitude shorter at ∼50 ms (16), despite the fact that PLCγ1 is primarily recruited to a single phosphotyrosine site on LAT (pY132), whereas Grb2 readily binds three phosphotyrosine sites (pY171, pY191, and pY226) (42). PLCγ1’s much slower dissociation kinetics likely contribute significant stabilization to the LAT condensate. This is consistent with observations in live T cells indicating that PLCγ1 recruitment is a rate-limiting step in the nucleation of LAT condensates (9, 50).

### PLCγ1 is rapidly activated at the membrane irrespective of LAT condensation state

We next extended the assay to measure the enzymatic activity of PLCγ1 hydrolyzing PIP_2_ into DAG (Figure 3A). PLCγ1 activity can be measured by tracking either PIP_2_ depletion (57) or DAG buildup (34) on the membrane (see example data in Figure S2A-B). Here we focus on DAG readout using a fluorescent DAG sensor engineered from the C1 domain of PKC theta. Membrane recruitment of the DAG sensor at equilibrium is linearly proportional to DAG membrane density (Figure S2C) and equilibration times were typically within 1 minute. PLCγ1 is recruited to the pLAT from lysate in its autoinhibited form and exhibits essentially no catalytic activity as shown in Figure 3B.

**Figure 3.**
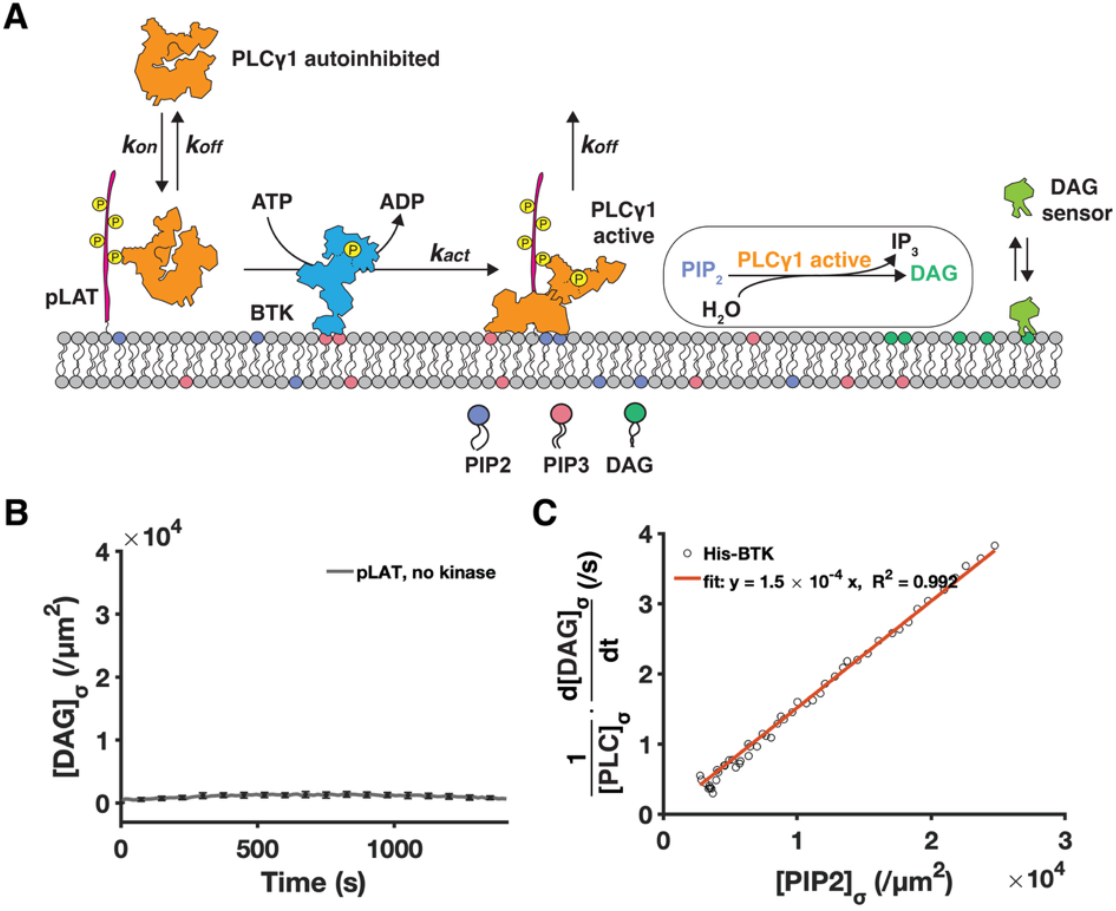
Reconstituted PLCγ1 activation assay. (A) Schematic of the reconstituted PLCγ1 activation assay. PLCγ1 is recruited to the membrane through prephosphorylated LAT and activated by the ITK surrogate BTK. Once phosphorylated at Y783, PLCγ1 catalytic activity is monitored using a fluorescence-based diacylglycerol (DAG) sensor. *k*_*on*_ is the rate of PLCγ1 membrane adsorption. *k*_*off*_ is the rate of PLCγ1 membrane dissociation. *k*_*act*_ is the rate of PLCγ1 phosphorylation and autoinhibition release. (B) Without membrane-bound kinase, PLCγ1 recruits to the pLAT in the autoinhibited conformation. pLAT was incubated at approximately 500 molecules/μm^2^, PLCγ1 was added at 15 nM, and DAG sensor was added at 200 nM. (C) The linear proportionality between PLCγ1 catalytic rate and substrate (PIP_2_) density, indicating that for active PLCγ1, *K*_*M*_ ≫ [*PIP*_2_]_σ_. (Membrane: 90:4:2:4 DOPC:Ni-NTA:PIP_2_:PIP_3_.)

In T cells, PLCγ1 activation is primarily mediated by the TEC family kinase ITK (48). Here, we employed Bruton’s tyrosine kinase (BTK), another TEC family kinase and the native activator of PLCγ2 in B cells (58, 59), as a surrogate for ITK for three reasons. First, BTK and ITK share the same domain organization and exhibit high sequence similarity, particularly within their kinase domains (60). Like ITK, BTK is recruited to the membrane via its PH-TH domain binding to PIP_3_ and can phosphorylate PLCγ1 (61-64). Second, full-length human ITK is challenging to purify due to its tendency to oligomerize and its persistent association with chaperonins (65, 66). However, full-length human BTK can be readily expressed and purified from E. coli (62). Third, the activation of ITK requires phosphorylation at its activation loop by SRC family kinase LCK (67), while BTK can dimerize and auto-phosphorylate on PIP_3_-containing membranes (68). The intrinsic ability of BTK to self-activate in the absence of upstream kinases offers a simplified—and more quantitative—system for the focused investigation of PLCγ1 activation, free from the influence of an ITK-activating kinase.

First, to avoid the added complexity of the PIP_3_-mediated BTK recruitment and activation, we created a system in which BTK was preactivated and stably tethered to the membrane using full-length BTK fused to an N-terminal His-SUMO tag (His-BTK) (Figure 4A). His-BTK was anchored to supported lipid bilayers via nickel-histidine interactions and subsequently activated by ATP-driven autophosphorylation over 30 minutes, yielding a membrane platform with catalytically active and stably associated His-BTK along with pLAT. Upon the addition of PLCγ1, we observed rapid DAG production, indicating robust activation of pLAT-recruited PLCγ1 by His-BTK on the membrane (Figure 4B).

**Figure 4.**
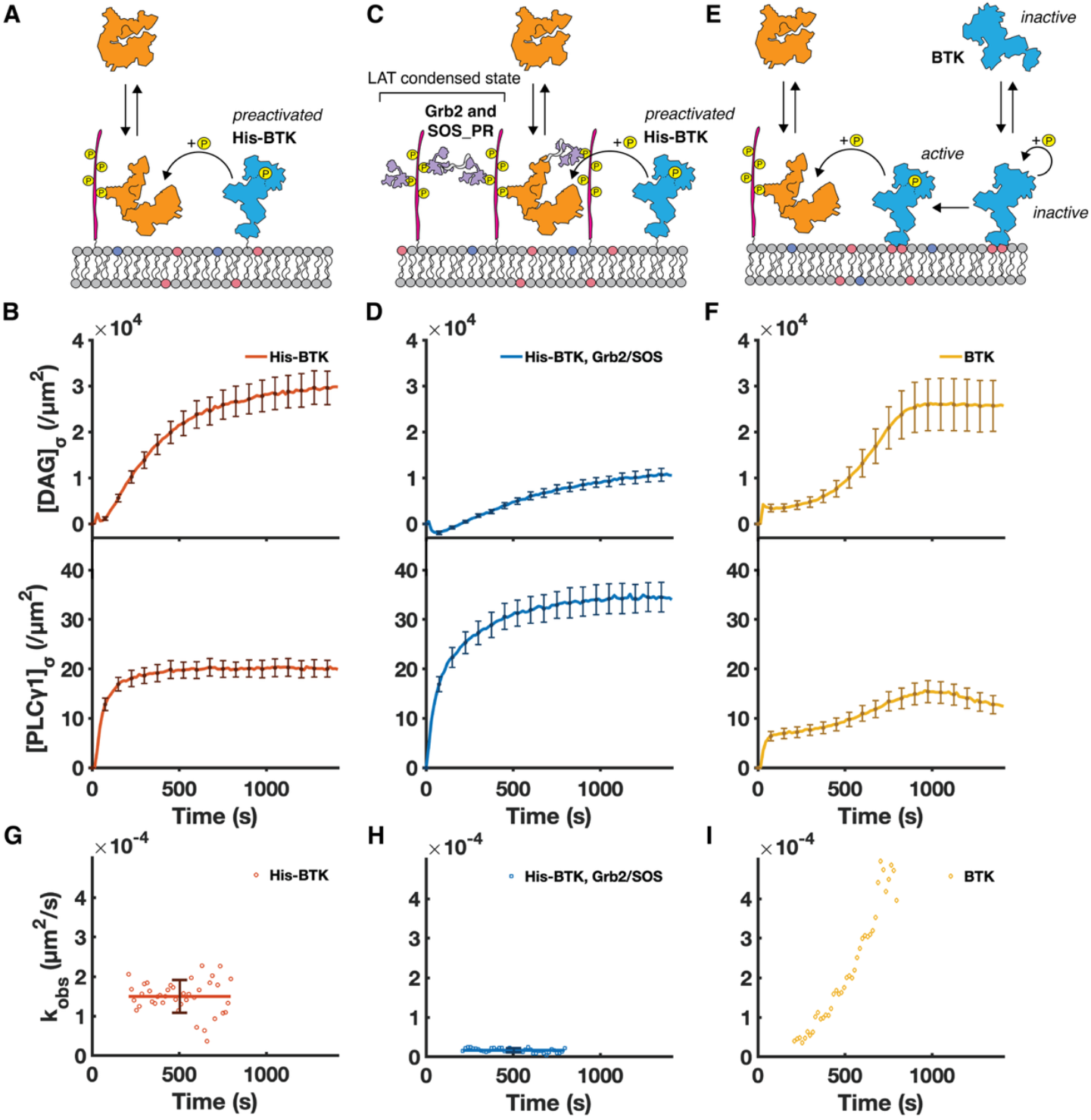
PLCγ1 is rapidly activated at the membrane irrespective of LAT condensation state. Schematic and kinetic traces of PLCγ1 activation under three conditions: (A-B) preactivated His-BTK and dispersed pLAT, (C-D) preactivated His-BTK and condensed pLAT, (E-F) native (initially autoinhibited) BTK and dispersed pLAT. (G-I) Measured (dotted line) and fitted (solid line) *k*_*obs*_ for each condition. PLCγ1 is rapidly activated with membrane-bound BTK irrespective of LAT condensation state. pLAT was incubated at approximately 500 molecules/μm^2^, PLCγ1 was added at 15 nM, and DAG sensor was added at 200 nM. For His-BTK condition, 50 nM of His-BTK was incubated on the membrane before the addition of the reaction mixture. For BTK condition, 50 nM of BTK was added along with the reaction mixture. (Membrane: 90:4:2:4 DOPC:Ni-NTA:PIP_2_:PIP_3_.)

The assay configuration described here enables quantitative characterization of PLCγ1 activity under several key physiological conditions. Most critically, the nSH2-mediated recruitment of PLCγ1 to membrane pLAT positions the enzyme at its physiological localization near the membrane, which substantially impacts its ability to bind its PIP_2_ substrate. Previous characterizations of PLCγ1 catalytic rate in reconstitution have been performed by directly incubating autoinhibited PLCγ1 with substrate in solution, micelles, or liposomes (55, 69-71) without nSH2-mediated membrane recruitment or phosphorylation-mediated autoinhibition release. Measurements from such assays cannot be quantitatively mapped to PLCγ1 activity in the membrane context. To interpret kinetic measurements of PLCγ1 in the experiments described here, we consider a membrane surface Michaelis-Menten mechanism for PIP_2_ hydrolysis to DAG:

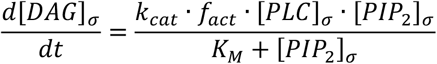

where *k*_*cat*_ and *K*_*M*_ are the standard Michaelis-Menten parameters and we use […]_σ_ to denote surface density. PLCγ1 activity levels are governed by its adsorption, activation, and desorption kinetics, and vary with experimental conditions (Figure 3A). We introduce the parameter *f*_*act*_ to account for the fraction of membrane-associated PLCγ1 molecules that have entered the active state. For experiments using preactivated His-BTK, *f*_*act*_ stays relatively constant after PLCγ1 membrane recruitment equilibrates (see SI for details). Measurements of PLCγ1 catalytic rate as a function of substrate (PIP_2_) density under steady state *f*_*act*_ exhibit nearly linear proportionality (Figure 3C), indicating *K*_*M*_ ≫ [*PIP*_2_]_σ_ over the full range of experimental conditions (see SI for details). Even under conditions where PLCγ1 and its substrate are both highly concentrated on the membrane, substrate binding remains rate-limiting for the catalytic reaction. This behavior contrasts with SOS, which operates in a catalytic rate-limited regime under the same membrane-localized conditions (29, 72). With this understanding of nSH2-recruited PLCγ1 behavior at the membrane, the rate equation simplifies to an essentially bimolecular reaction with apparent rate constant, *k*_*obs*_ ≈ (*k*_*cat*_ · *f*_*act*_)/*K*_*M*_:

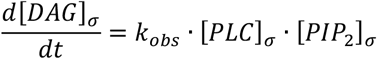

We utilize this kinetic description to extract values of *k*_*obs*_ from experimental measurements.

PLCγ1 catalytic rates are compared under three different situations: preactivated His-BTK and dispersed pLAT (Figure 4A-B), preactivated His-BTK and condensed pLAT (Figure 4C-D), and native (initially autoinhibited) BTK and dispersed pLAT (Figure 4E-F). The condensation state of LAT was controlled by including Grb2 and the proline-rich region of SOS (SOS_PR_) together with PLCγ1 during the assay. At Grb2 and SOS_PR_ concentrations of 5 μM and 2 μM, respectively, robust LAT:Grb2:SOS condensation occurred (16, 34) as shown in Figure S3. For preactivated His-BTK and either dispersed or condensed pLAT, we observed relatively constant values of *k*_*obs*_ after about 200s, when PLCγ1 adsorption and activation levels have reached steady-state. Consistent with earlier observations (34), we measure PLCγ1 to achieve its maximal activity on dispersed pLAT (*k*_*obs*_ = 1.5 × 10^−4^ *μm*^2^/*s*, Figure 4G). Moderately increased PLCγ1 recruitment was measured with condensed pLAT (likely reflecting avidity effects) but the per-molecule PLCγ1 activity was distinctly lower (*k*_*obs*_ = 1.6 × 10^−5^ *μm*^2^/*s*, Figure 4H). The reduced activity seen from condensed pLAT may reflect restricted substrate access within the reconstituted condensate. However, we note that boundary-to-interior ratios of physiological condensates are much larger suggesting that peripheral PLCγ1 (likely less restricted) may be the primary contributor to signaling in cells (9). In any case, PLCγ1 clearly does not require the condensed state of LAT—contrasting SOS. We also observed that PIP_3_ moderately enhances PLCγ1 catalytic activity, but it is not required (Figure S5A).

Observations of PLCγ1 activation using full-length native human BTK reveal distinctly different behavior. Native BTK is recruited to the membrane by PIP_3_ binding and arrives initially in an autoinhibited state (62). Once at the membrane, BTK slowly autophosphorylates and ultimately begins to activate PLCγ1 (Figure 4E-F). In this case with wild-type BTK, *k_obs_* did not reach a steady state over the course of the reaction (Figure 4I), reflecting the slow BTK autoactivation process in this experimental configuration and an increasing fraction of activated PLCγ1 (*f*_*act*_) over time. We note the overall *k*_*obs*_ value exceeded that measured with His-BTK, potentially reflecting differences in intrinsic catalytic efficiency between wild-type BTK and the His-SUMO-tagged construct. Consistent with literature (62, 63), fluorescently labeled BTK was only recruited to the membrane and activated in the presence of PIP_3_ (Figure S5B-C).

### The ITK surrogate BTK cross-phosphorylates LAT in addition to PLCγ1

An intriguing observation emerged when examining PLCγ1 membrane recruitment kinetics in the context of full-length BTK (Figure 4F). PLCγ1 recruitment no longer followed a simple binding curve. Instead, an initial wave of membrane association occurred early, followed by a second, delayed wave that coincided with the onset of DAG production (Figure 4F). The tight time correlation suggests that the second wave of recruitment is related to BTK activation. Two potential mechanisms could give rise to this observation: one possibility is that activated BTK directly recruits PLCγ1 to the membrane; alternatively, BTK may phosphorylate more LAT at Y132, generating more recruitment sites for PLCγ1.

We first evaluated the possibility that activated BTK directly recruits PLCγ1 to the membrane. We simultaneously added BTK and PLCγ1 to supported lipid bilayers containing 4% PIP_3_ (Figure 5A). Although BTK was recruited to the membrane, no detectable PLCγ1 recruitment was observed (Figure 5B), indicating that direct binding to BTK does not mediate additional PLCγ1 membrane association.

**Figure 5.**
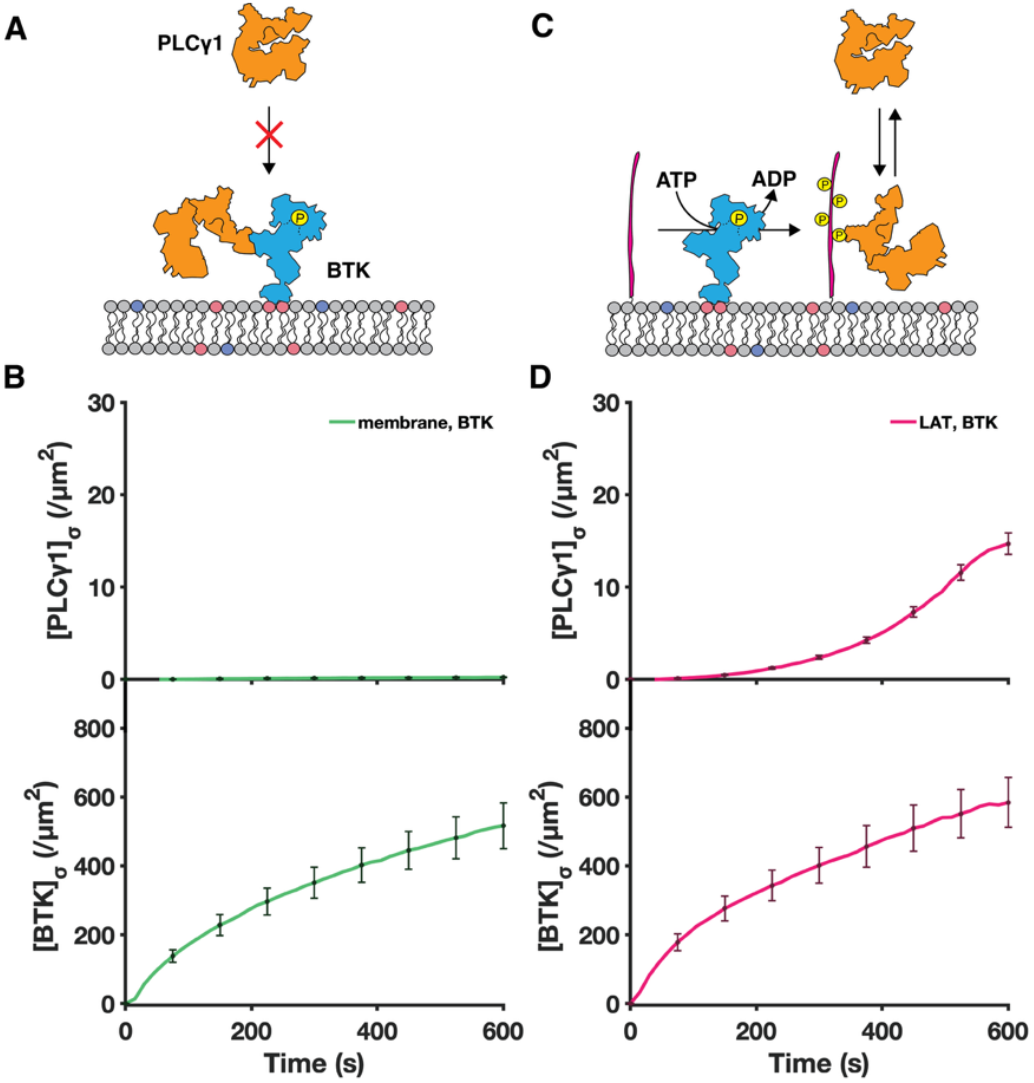
The ITK surrogate BTK cross-phosphorylates LAT in addition to PLCγ1. The second phase of PLCγ1 recruitment may arise from two mechanisms: direct BTK-PLCγ1 recruitment or BTK-mediated LAT phosphorylation, which in turn recruits additional PLCγ1. (A) Schematic of BTK directly recruiting PLCγ1. (B) Although BTK is robustly membrane-recruited, PLCγ1 is not recruited under this condition. 50 nM of BTK-555 was added along with 15 nM of PLCγ1. (C) Schematic of BTK phosphorylating LAT and thereby enabling additional PLCγ1 recruitment. (D) BTK recruits to the membrane and leads to a delayed PLCγ1 recruitment. LAT was incubated at approximately 500 molecules/μm^2^. (Membrane: 90:4:2:4 DOPC:Ni-NTA:PIP_2_:PIP_3_.)

Next, to examine the possibility that BTK can cross-phosphorylate LAT, we preincubated unphosphorylated LAT on supported lipid bilayers containing 4% PIP_3_ and added BTK and PLCγ1 (Figure 5C). We observed PLCγ1 recruitment to the membrane to begin after a delay of approximately 200 seconds, reminiscent of the timing of the second recruitment wave observed previously (Figure 5D). With no other kinase in the system, BTK is the sole source of both LAT phosphorylation and PLCγ1 recruitment.

Observation of LAT phosphorylation by BTK provides strong evidence that ITK would similarly phosphorylate LAT in physiological LAT condensates. The kinase domain of BTK and ITK have similar substrate specificity (60, 73). ITK has been reported to exhibit a modestly slower catalytic rate compared with BTK (using a synthetic tyrosine kinase substrate optimized for LCK activity) due to differences in their activation loop sequences (60). Although substitution of the BTK activation loop into ITK does not substantially alter the *K*_*m*_, it increases the *k*_*cat*_ from 0.54 min^-1^ to 1.75 min^-1^, indicating that the native ITK activation loop reduces catalytic turnover by approximately fourfold relative to BTK (60). Our measurements indicate that BTK and ZAP70 have similar per-molecule *in situ* catalytic rates for LAT Y132 phosphorylation on the membrane (details in SI, Figure S6). Additionally, an orthogonal study on the substrate specificity of the human tyrosine kinome (73) ranked BTK with a higher specificity towards LAT Y132 than ZAP70 (details in SI). We note that the Y132 (PLCγ1 binding site) is a relatively weak substrate for ZAP70 compared with the Grb2 sites (Y171, Y191, and Y226) (5, 74). Unlike ZAP70, which is stoichiometrically linked to TCR, ITK is stoichiometrically linked with LAT itself through Gads and SLP76 (75). For physiological LAT condensates observed in primary T cells, LAT molecules outnumber ZAP70 molecules by roughly 100-fold (9, 76). Thus, ITK could vastly outnumber ZAP70 in the condensate. Additionally, LAT condensates have been observed to continue growing in T cells even after the initiating pMHC:TCR complex, and presumably ZAP70 along with it, is gone (9)—suggesting there may be another source of LAT phosphorylation once the condensate is nucleated.

### Cross-phosphorylation of LAT by ITK creates positive feedback for LAT condensate growth

We examine the potential consequences of an ITK-driven positive feedback mechanism of LAT phosphorylation in condensates with computational simulations. For simplicity, we employ a spatially homogeneous (well-mixed) model to explore the relationship between LAT phosphorylation levels and kinase-phosphatase balance. While this analysis does not explore spatial reaction-diffusion aspects of condensate growth, it does reflect the net direction of the phosphorylation-dephosphorylation reactions with informative results. In this model, the kinase ZAP70 phosphorylates LAT with a rate constant *β*, and phosphatases such as CD45 and SHP-1 dephosphorylate LAT with a rate constant *ξ*. Once recruited to the LAT condensate, ITK phosphorylates additional LAT molecules with a rate constant *α* (Figure 6A). While ITK (through its associated adaptor proteins) is bound to phosphorylated LAT, phosphatases are unable to access and dephosphorylate the occupied phosphotyrosine sites.

**Figure 6.**
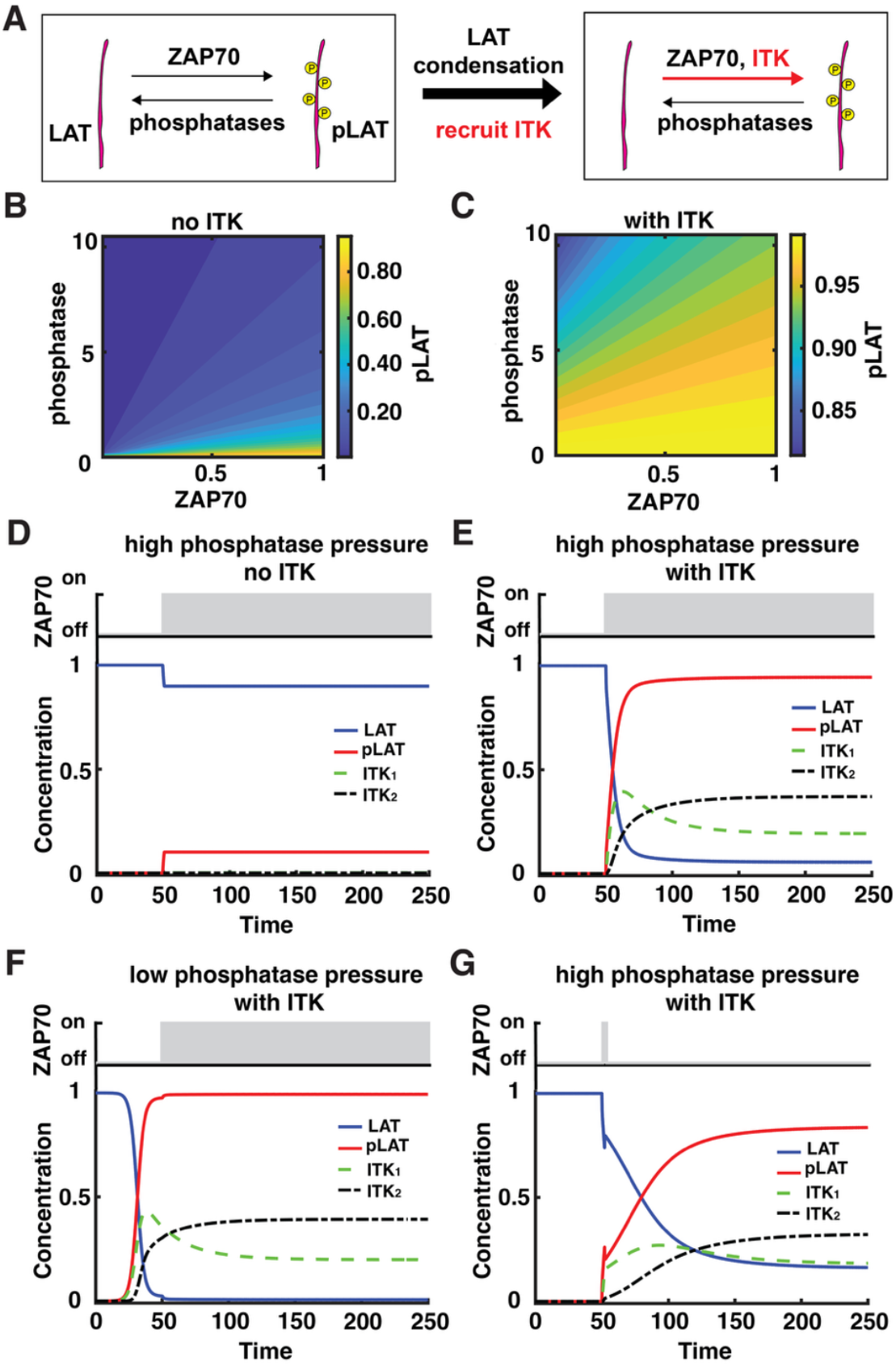
Model simulations reveal ITK-driven positive feedback leading to bistability in LAT phosphorylation. (A) Schematic showing that LAT condensates recruit ITK and shift the balance between kinase and phosphatase activities on LAT. (B-C) Steady-state levels of pLAT from phosphatase and ZAP70 titration in the absence (B) or presence (C) of ITK. (D-G) For kinetic simulations, phosphatase and ITK are present from t = 0, and ZAP70 is activated at t = 50. (D) Under high phosphatase pressure and no ITK, pLAT remains low. (E) Under high phosphatase pressure with ITK, pLAT can rapidly switch to a high-phosphorylation state. (F) Under low phosphatase pressure with ITK, the system becomes unstable and reaches a high pLAT level before ITK onset. (G) Under high phosphatase pressure with ITK, the system still reaches a high pLAT state even after ZAP70 is turned off at t = 52. All units are non-dimensionalized according to methods.

In T cells, ITK associates with LAT primarily through the adaptor proteins SLP76 and Gads, which interact with phosphorylated tyrosine sites on LAT as well as with PLCγ1. Together with Grb2 and SOS, all of these proteins—including ITK—collectively define the LAT condensate through a percolating bond network of protein-protein interactions (14, 75, 77, 78). In this model, ITK recruitment to the LAT condensate is represented as a simplified two-step process. We defined the initial association and dissociation rate constants of Gads-SLP76-ITK binding to pLAT (*ITK*_1_) by *μ*_1_ and γ_1_, respectively. Further crosslinking between ITK and LAT in the condensate leads to a second state of complex formation (*ITK*_2_) that occurs with binding and unbinding rate constants *μ*_2_ and γ_2_, respectively. Use of two distinct association states is a minimalist approach to capturing the experimentally-observed avidity effects, which substantially alter dissociation kinetics of proteins in the condensed state (16, 29). We then proposed the following rate equations to describe the levels of phosphorylated LAT and pLAT-associated ITK complexes:

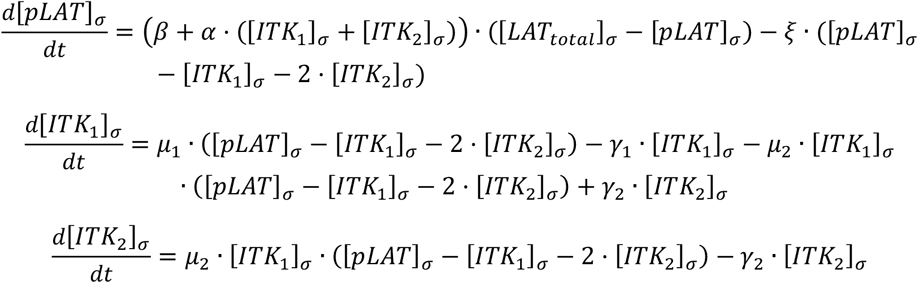

After non-dimensionalizing the concentrations and kinetic parameters, we first examined a system lacking ITK-mediated positive feedback by imposing a high dissociation rate for ITK (γ_1_ ≫ 1, γ_2_ ≫ 1), thereby minimizing its contribution to LAT phosphorylation. In this case, the steady-state level of phosphorylated LAT is determined primarily by the balance between ZAP70-mediated phosphorylation and background dephosphorylation by phosphatases (Figure 6B). As expected, the addition of ITK shifts the balance toward a higher steady-state level of pLAT (Figure 6C). To gain further insight into the system’s dynamics, we analyzed the kinetics of LAT phosphorylation following ZAP70 activation, which was initiated at *t* = 50.

Under high phosphatase pressure, and after ZAP70 activation, the system reaches a steady state characterized by a low level of phosphorylated LAT (Figure 6D). This corresponds to a situation in which the active ZAP70 on a triggered TCR is insufficient to overcome background phosphatase activity. Upon introduction of ITK into the system, the same ZAP70 input drives pLAT to a high steady state (Figure 6E). At low phosphatase pressure, the system becomes unstable and small fluctuations (noise) can drive it toward a high pLAT state even before ZAP70 activation (Figure 6F). This result further demonstrates that a high level of phosphatase activity in T cells is necessary to prevent spontaneous LAT phosphorylation and T cell activation in the absence of antigen stimulation. We also observed that even when ZAP70 activity was turned off shortly after activation (at *t* = 52), ITK kinase activity alone was sufficient to drive and sustain the transition to the high-pLAT state (Figure 6G). This result is consistent with experimental observations in T cells where LAT condensate growth can be observed even after dissociation of the initiating agonist:TCR complex and its associated ZAP70 activity (9).

## Discussion

PLCγ1 and SOS are both critical downstream signaling molecules recruited to and activated in LAT condensates during TCR signaling. However, the mechanisms by which LAT condensation regulates these two molecules differ dramatically. While the condensed state of LAT strongly promotes SOS activation (via a type of kinetic proofreading process) (29), PLCγ1 shows no preference for the condensed state and is robustly active even from the dispersed LAT state. The indifference of PLCγ1 to the LAT condensation state can be rationalized, in part, by its strong binding to the pY132 site (here measured mean dwell time ∼100 s). SOS interactions with dispersed pLAT, by contrast, reflect monovalent binding kinetics of the bridging Grb2 molecules with mean dwell times orders of magnitude shorter around ∼50 ms (16). The likelihood of *in situ* cross-phosphorylation of LAT Y132 by the PLCγ1 activator, ITK, suggested a different way in which LAT condensation may indirectly control PLCγ1 activation.

ITK-mediated LAT cross-phosphorylation in condensates would create a positive feedback loop that, in concert with strong phosphatase activity, can establish a bimodality in LAT phosphorylation levels (Figure 7). One consequence of this bimodality is that cellular phosphatase activity can be maintained at a relatively high level, assuring that any spurious LAT phosphorylation is rapidly reversed to maintain a very low background level of pLAT (as experimentally observed) (6). The high phosphatase activity against dispersed pLAT establishes a resistance, or negative gain step, after TCR activation that gates LAT phosphorylation. Once nucleated, however, positive feedback by ITK phosphorylation of LAT provides signal amplification beyond the limited numbers of ZAP70 molecules. We note that phosphatase exclusion in LAT condensates has been identified as another form of positive feedback (9, 15), but this is a double negative effect that still relies on ZAP70 phosphorylation and therefore has far more limited potential than ITK. While both mechanisms may contribute, the fact that ITK copy numbers are proportional to LAT, rather than TCR, suggests it is likely the dominant force.

**Figure 7.**
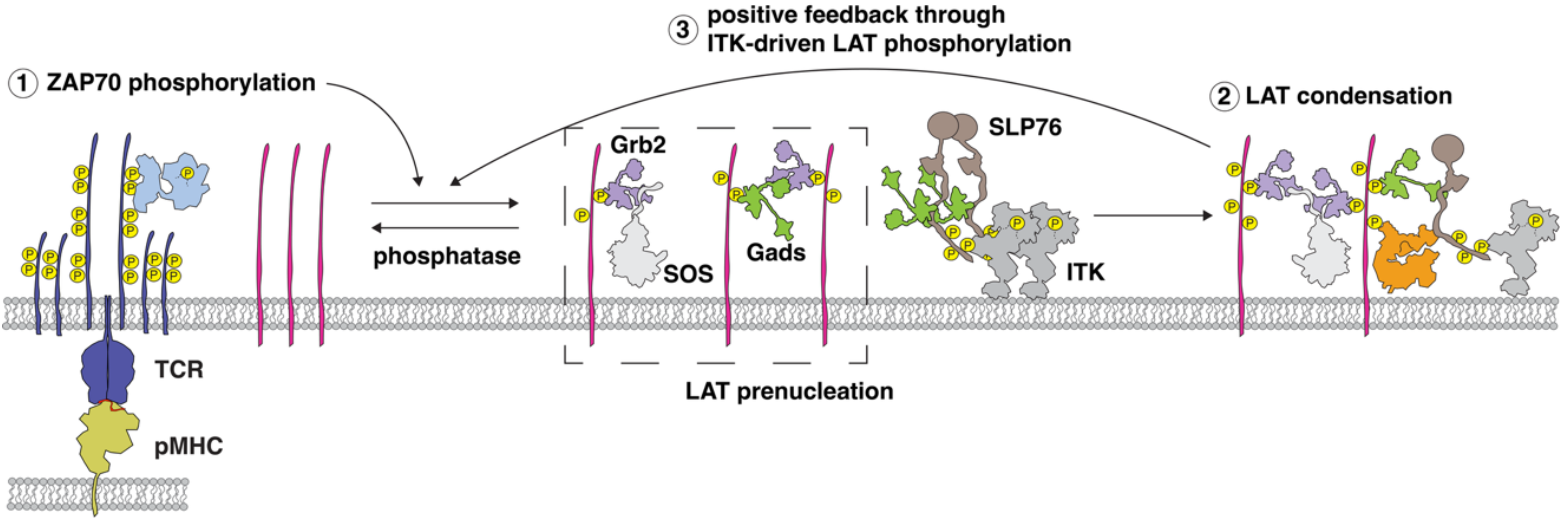
Proposed mechanism of ITK-driven positive feedback. Upon TCR stimulation and ZAP70 activation, a fraction of LAT molecules proximal to the pMHC-TCR engagement site becomes phosphorylated. Some pLAT molecules bind adaptor proteins and form prenucleation clusters, while others are rapidly dephosphorylated under high phosphatase pressure. A subset of prenucleated LAT clusters recruits PLCγ1 and ITK, stabilizing and driving rapid growth to form mature, signaling-competent LAT condensates. Within condensates, pLAT is protected from phosphatase activity, and ITK-mediated positive feedback further reinforces LAT phosphorylation, resulting in bimodality in LAT phosphorylation—where LAT is phosphorylated within condensates but remains largely unphosphorylated outside. Consequently, PLCγ1 activation is confined to LAT condensates.

Functionally, bimodality in pLAT levels would ensure that nonspecific LAT phosphorylation events (noise) remain suppressed while true, sustained, antigen signals can be strongly amplified. Since, as observed here, PLCγ1 recruitment depends entirely on pLAT, a stable steady state with very low pLAT levels keeps PLCγ1 safely autoinhibited in the cytosol even with occasional spurious LAT phosphorylation events (79, 80). However, once a true agonist:TCR complex has nucleated a LAT condensate, strong positive feedback and a ZAP70-independent growth mechanism through ITK assure strong PLCγ1 recruitment and activation, leading to a definitive cellular calcium response, as experimentally observed by Morita et al (20).

An intriguing observation from our computational studies is that while ZAP70 is required to nucleate the LAT condensate, it is not required for sustained growth. This result is consistent with several experimental observations in primary T cells. First, LAT condensates can be observed to continue to grow even after the initiating pMHC:TCR complex has dissociated (9, 20). Second, there is no correlation between the size of a LAT condensate and the duration of the pMHC:TCR complex that initiated it (9). Once nucleated, condensates evidently grow (and reach a limited size) independently of pMHC:TCR binding and ZAP70 dwell time. Taken together, these computational results and live cell observations suggest that TCR-activated ZAP70 provides a spark, which is necessary to nucleate LAT condensation via a type of ignition process. Once ignited, the condensates grow via ITK-driven phosphorylation of LAT independently from engaged TCR. While the extent of ITK-LAT phosphorylation under physiological conditions remains to be quantified, this spark-ignition hypothesis is consistent with all current experimental observations. More broadly, these results underscore how kinase substrate specificity is not a fixed parameter. Strong colocalization to a membrane surface or in a condensate can enable cross reactions, which are unlikely in the solution state, to occur—expanding the potential roles for specific kinases and phosphatases in signal propagation and attenuation.

## Acknowledgments

We thank the members of the Groves Laboratory for helpful discussion. This work was supported by the Novo Nordisk Foundation Challenge Program under the Center for Geometrically Engineered Cellular Systems and NIH Grant PO1 A1091580.

